# Genetic variation in *Eif2ak3* alters astrocyte-dependent stress tolerance in neurons

**DOI:** 10.64898/2026.09.29.755429

**Authors:** Shivesh Ghura, Elena Alvarez-Periel, Sarah E. Bond Newton, Cagla Akay-Espinoza, Kelly L. Jordan-Sciutto

**Author notes:** Corresponding author: Kelly L. Jordan-Sciutto University of Pennsylvania, 240 S. 40th St, Room 312 Levy Philadelphia, PA 19104-6030.

## Abstract

Subtle variations in the activity and signaling of PERK, an endoplasmic reticulum kinase, can shift stress response outcomes. PERK has three commonly occurring exonic single-nucleotide variants frequently inherited together as part of the PERK haplotype B (PERK-B), which occurs in 28% of the global population and is associated with several chronic neurodegenerative diseases, although its impact on PERK activity remains controversial. We hypothesized that PERK-B was associated with increased activity, altering stress response and stress tolerance, and that minorly but consistently elevated PERK activity, in combination with additional environmental or genetic risk factors, would lead to adverse outcomes. Comparison of stress response between PERK-A, the major PERK haplotype, and PERK-B in neuronal cultures using novel knock-in PERK-B mice alongside wild-type PERK-A mice revealed differential, transient changes. Following ER stress, astrocyte-containing neuronal cultures, but not neuron-only cultures expressing PERK-B exhibited similar levels of eIF2α phosphorylation, the canonical PERK substrate at PERK peak activation point (2h), but showed higher levels at 4–6 h, consistent with altered kinetics of PERK signaling. PERK-B neuronal cultures pretreated with a low-dose PERK-specific activator exhibited increased vulnerability to N-methyl-D-aspartate-induced death compared to PERK-A neuronal cultures, indicating impaired acquired stress tolerance. Additionally, neuron-astrocyte coculture experiments revealed that the observed difference in acquired stress tolerance was dependent on the astrocyte genotype. The association of PERK-B with transient changes in stress response and reduced stress tolerance with higher vulnerability to cellular stress, albeit transient and small/moderate in size, has clinical relevance based on its potential contribution to the overall genetic vulnerability in neurodegenerative diseases.

## Introduction

Common contributors to neuronal damage and dysfunction in many neurodegenerative diseases, such as immune dysregulation, neuroinflammation, and oxidative stress, are primarily driven by astrocytes, macrophages, and microglia^1–5^. In these cells, activation of multiple stress response pathways (e.g., antioxidant response, unfolded protein response [UPR]), orchestrated by multiple organelles (e.g., endoplasmic reticulum [ER], mitochondria), aims to restore homeostasis and, in cases of failure, leads to neuronal damage/dysfunction^6–8^.

Among the stress-related pathways induced by neurodegenerative insults is the integrated stress response (ISR), a ubiquitous, multipronged pathway that serves as a central node in adaptive signaling downstream of specific stressors, helping resolve stress and restore homeostasis^9,10^. The ISR is a framework of four kinases that converge on the phosphorylation of eukaryotic initiation factor 2α (eIF2α), which leads to a global pause in cap-dependent protein translation coupled with the selective translation of a subset of mRNAs, leading to adaptive proteome remodeling^11^. Activating transcription factor (*ATF4*) is one such mRNA^12^. ATF4 target genes, such as the ER chaperone binding protein (BiP), are involved in protein folding, nutrient uptake, and response to oxidative stress. During prolonged stress, ATF4 also induces the expression of CCAAT-enhancer-binding homologous protein (CHOP), which induces apoptosis^13^. As a transcription factor itself, CHOP induces the expression of growth arrest and DNA-damage inducible protein 34 (GADD34), which forms a complex with protein phosphatase 1c to dephosphorylate eIF2α, thereby providing negative feedback for stress-induced gene expression^14^. Importantly, depending on the strength and duration of stress, ISR activation can lead to stress resolution and restoration of cellular homeostasis or to cell death^9,15–20^.

Among the four ISR kinases, protein kinase R-like ER kinase (PERK) is triggered by the accumulation of unfolded/misfolded proteins in the ER or excessive Ca^2+^ efflux from the ER^21^. Specifically, BiP, which basally binds PERK and aids in keeping it inactive, is titrated away by such accumulation^22–25^. BiP and other protein chaperones are also calcium-regulated, facilitating stress sensing and activation in concert with stress signaling related calcium efflux^26^. Canonically, PERK aims to restore ER homeostasis via transcriptional upregulation of ATF4-target genes and downstream PERK targets, including *NFE2L2*, chaperones (e.g., *BIP*, *HSP70*), autophagy-related genes (e.g., *ATG5*, *P62*), and genes involved in antioxidant response (e.g., *HO-1*)^8,27^. Following stress resolution, BiP rebinds to the PERK luminal domain, inhibiting its oligomerization necessary for kinase activity^28,29^.

In the CNS, PERK signaling and downstream eIF2α phosphorylation act on local translation, regulating long-term potentiation and synaptic plasticity^30–34^. For example, alterations in the levels of synaptic proteins, such as PSD95 and CREB, secondary to altered eIF2α phosphorylation, are observed in models of traumatic brain injury^35^. Although genetic PERK knockdown has minimal impact on memory, subtle changes in specific behavioral tests reported in rodents suggest modulatory roles for PERK in neurons^30,36,37^. PERK also mediates ER stress and downstream inflammatory pathways in astrocytes^38–42^. These findings suggest that PERK regulation and function may have unique consequences in CNS cells.

Beyond the primary outcomes of PERK signaling, accumulating evidence also reveals that PERK activation preconditions cells to other stressors, such as subsequent exposure to excitotoxicity following ER stress^43–45^. Low/mild ER stress can trigger an adaptive response and is beneficial for the survival of cells challenged with a subsequent stressor, a phenomenon termed hormesis or stress tolerance^46–50^. This added level of PERK functionality highlights another dimension to PERK’s contribution to stress response.

Given its critical role in cellular homeostasis, it is not surprising that PERK signaling is dysregulated in a range of neurodegenerative conditions. Altered PERK signaling has been reported in Alzheimer’s disease (AD)^51^, Parkinson’s disease, progressive supranuclear palsy^52,53^, and HIV-associated neurocognitive disorders (HAND)^54^, among others. However, the complex crosstalk of PERK signaling with other ISR kinases and stress pathways hinders efforts to develop therapeutic approaches targeting PERK^55,56^, as exemplified by the demonstrated benefit of both PERK inhibition^35,57–72^ and activation^73–81^ in models of neurodegeneration^82^.

In addition to complex regulatory mechanisms, PERK has substantial variability in its genetic locus in humans. *EIF2AK3*, the gene encoding PERK, has commonly occurring single-nucleotide variants (SNVs). Among these, one *EIF2AK3* promoter SNV (rs6547787), along with three nonsynonymous exonic *EIF2AK3* SNVs (rs13045, rs867529, and rs1805165) form the PERK B haplotype, which occurs in approximately 28% of the global population, compared with the prevalent PERK A haplotype, which occurs in 64%^82^. We recently developed triple knock-in mice expressing the 3 exonic SNVs (hereto “PERK-B”) reported that recombinant PERK-B exhibits higher kinase activity than recombinant PERK-A^83^, which has several implications for neurodegeneration, given the reported association of PERK-B-related SNVs with increased risk in several neurodegenerative disorders^51,52,54,84,85^. However, it remains unclear how the PERK-B haplotype might be contributing to disease processes. PERK-B was more active in lymphocytes in one study^82^, yet functioned as a hypomorph in fibroblasts in another^86^. Therefore, understanding the contribution of PERK genetic variation to neurodegeneration requires elucidating its impact on PERK signaling in the CNS.

Here, we aimed to discern the specific impact of the PERK-B haplotype on stress response and stress tolerance in distinct CNS cell types by comparing neuronal, neuron-only, astrocytic, and co-cultures between the novel triple knock-in PERK-B/B mice and the wild-type PERK-A/A mice. We hypothesized that PERK-B would have increased kinase activity, augmenting the stress response while diminishing stress tolerance, an effect that would be cell type-specific. Our findings suggest that the subtle alterations induced by these variants, despite their relatively high frequency in the general population and moderate effect size, might only manifest in specific stress contexts or paradigms.

## Materials and Methods

### Animals

The triple knock-in PERK-B/B mice were generated as previously described^83^, by introducing three single-point mutations (S132C, R162Q, and S700A) into the *Eif2ak3* locus in C57BL/6 mice (i.e., PERK-A/A). All animal protocols were approved by the Institutional Animal Care and Use Committee at the University of Pennsylvania (protocol no. 806053) and conducted in accordance with the NIH guidelines, complying with ARRIVE guidelines for animal care and animal experimentation (https://arriveguidelines.org). Mice were deeply anesthetized with CO_2_ in a chamber filled with 100% CO_2_ at a fill rate of 30%–70% displacement/min. After observing breathing cessation for at least 2 min, euthanasia was confirmed by cervical dislocation. For all experiments, homozygous PERK-AA and PERK-B/B mice were used.

### Isolation and culture of primary neuronal, astrocytic, and neuron-astrocyte cocultures

Primary mouse cortical neuronal cultures were prepared from embryonic day 16–18 PERK-A/A or PERK-B/B mouse embryos, as previously described^87–89^. Briefly, cortices from embryos were isolated and incubated in 0.5 mL trypsin (0.025%) and 4.5 mL HBSS for 15 min at 37°C, after which 5 mL HBSS was added for another 5-min incubation. After the addition of 5 mL fresh HBSS, the tissue was incubated for an additional 5 min at 37°C. Fetal bovine serum (FBS)-containing medium was added to cells, which were triturated and counted. Cells were resuspended in neurobasal medium supplemented with 1% penicillin/streptomycin, 2% B27, and 0.5% GlutaMAX and plated on poly-l-lysine-coated 6- or 24-well plates (USA Scientific, Ocala, FL) at a concentration of 300,000 cells/mL. Cultures were maintained at 37°C in 5% CO_2_ for up to 21 days with half-medium changes every 5–6 days and treated on days in vitro (DIV) 16–18. Of note, pure neuronal cultures were obtained by treating cultures with 2 µM arabinofuranosylcytosine (AraC) on DIV 2–3, whereas neuronal cultures were not treated with AraC. Primary *astrocyte cultures* were prepared from post-natal day 0–2-day-old PERK-A/A or PERK-B/B mouse pups, as previously described^90^. Briefly, cortices from pups were isolated, incubated in 0.5% trypsin-EDTA for 5 min at 37°C, resuspended in FBS-containing medium, and centrifuged at 600 *g* for 2 min. The pelleted cells were resuspended and passed through 100-µm and 40-µm cell strainers, followed by incubation in Dulbecco’s modified Eagle medium supplemented with 1% penicillin/streptomycin, 10% FBS, and 0.5% GlutaMAX. Cells were grown in 75-cm^2^ tissue culture flasks until confluency, with medium changed every 3–4 days and passaged once. Once confluent, cultures were plated in 6- or 24-well plates at 140,000 cells/well. Treatments were performed once the cultures reached confluency. For *neuron-astrocyte cocultures*, enriched neurons and astrocytes were isolated separately. Briefly, astrocytes were purified using a shake-off protocol, as described^91^, and subsequently added to plates containing pure neurons on DIV 16–20. The neuron/astrocyte ratio was 2.5:1. Next, 24–48 h after the addition of astrocytes, the neuron-astrocyte cocultures were treated with AraC to arrest astrocyte proliferation. The coculture medium was neurobasal medium supplemented with 1% penicillin/streptomycin, 2% B27, 0.5% GlutaMAX, and the G5 supplement, to ensure astrocyte survival.

### Chemicals and reagents

Antibodies to the following proteins were used: PERK (clone C33E10; catalog #3192), eIF2α (clone L57A5; catalog #2103), phospho-eIF2α (p-eIF2α; clone D9G8; catalog #3398), beta-actin (clone 8H10D10; catalog #3700), CHOP (clone L63F7; catalog #2895), and GFAP (clone GA5, catalog #3670) from Cell Signaling Technology (Danvers, MA); BiP (catalog #610978) from BD Transduction Laboratories (San Jose, CA); MAP2 (catalog #ab5392) from Abcam (Cambridge, UK); MAP2 (catalog #801801) from BioLegend (San Diego, CA); GFAP (catalog #Z033429-2) from Dako (Glostrup, Denmark); p-eIF2α (catalog #44-728G) from Invitrogen (Waltham, MA);; goat-anti rabbit FITC (catalog #111-095-144), goat anti-mouse Cy5 (catalog #115-175-146), and goat-anti chicken Cy3 (catalog #103-165-155) from Jackson ImmunoResearch (West Grove, PA). The following chemical reagents were used: RIPA buffer (catalog #9806, Cell Signaling Technology), N-methyl-D-aspartate (NMDA; catalog #0114/50; Tocris, Bristol, UK), anhydrous dimethyl sulfoxide (DMSO; catalog #276855; Sigma Aldrich, St. Louis, MO), DAPI (catalog #D21490, Thermo Fisher Scientific, Waltham, MA), thapsigargin (catalog #11381; Tocris), tunicamycin (catalog #351610, Thermo Fisher Scientific), and the PERK activator CCT020312 (PA; catalog #324879; Millipore Sigma, Burlington, MA).

### Immunoblotting

Tissue and cell lysates for immunoblotting were prepared as previously described^87^. Briefly, mouse tissues from specific brain regions were homogenized and solubilized in ice-cold RIPA buffer supplemented with 1:100 Halt^®^ protease and phosphatase inhibitor (Thermo Fisher Scientific) using a Dounce homogenizer. Extracts were then sonicated thrice at 7-sec intervals, with 10-sec breaks, while on ice. The lysates were centrifuged at 12,000 *g* at 4°C for 20 min. Cell culture samples were prepared by rinsing cells twice with phosphate-buffered saline (PBS), followed by lysis using ice-cold RIPA buffer supplemented with 1:100 Halt^®^ protease and phosphatase inhibitor. Protein supernatants were collected with centrifugation at 20,000 × *g* for 10 min at 4°C.

Protein concentrations were determined using a BCA protein assay kit (Thermo Fisher Scientific). Next, 10–20 μg protein from each tissue sample or 5–10 μg total protein from each cell culture sample was loaded onto 4%–12% Bis-Tris NuPAGE Novex gels (Thermo Fisher Scientific). After separation, the proteins were transferred to PVDF membranes, which were blocked with Tris-buffered saline plus 0.1% Tween 20 (TBS-T) supplemented with 5% bovine serum albumin (BSA; MilliporeSigma) for 1 h at room temperature. The membranes were incubated with indicated primary antibodies overnight at 4°C. After three washes in TBS-T, the membranes were incubated with appropriate horseradish peroxidase-conjugated secondary antibodies (1:1000 in TBS-T with 5% BSA) for 30 min at room temperature. Bands were visualized by chemiluminescence using Luminata Classico ECL (MilliporeSigma), and images were captured using film or the ChemiDoc Touch imaging system (Bio-Rad, Hercules, CA). Equal loading and even transfer of samples were confirmed by probing for beta-actin protein. Densitometric analysis of band intensities was conducted using Bio-Rad’s ImageLab software (version 6.0), and all bands were normalized to beta-actin.

### Acquired stress tolerance paradigm

Neuronal cultures and neuron-astrocyte cocultures were plated in 24-wells with 500 µL medium/well, as described above. At the time of experiments, cultures were pretreated with the PERK activator PA (0.05, 0.1, or 0.5 µM) or with tunicamycin (10 µg/mL) for 1 h, followed by a 5-h recovery consisting of two medium changes to dilute the stressors and to prevent additional stress or neurotoxicity due to media changes. First, 250 µL of medium was changed. Next, 250 µL of medium was removed, and 500 µL of fresh medium was added (67% media change). Subsequently, the cultures were treated with 10 µM NMDA for 20 h.

### Immunostaining

Immunofluorescence staining of cultures was performed as previously described^87^. Briefly, after treatments, cultures were washed with PBS and fixed with 4% paraformaldehyde for 20 min. After two washes with PBS and three washes with PBS supplemented with 0.1% Tween20 (PBS-T), cultures were incubated with the blocking/permeabilization solution (0.2% BSA and 0.1% Triton-X in PBS) for 60 min. After three washes with PBS-T, cultures were incubated with primary antibodies against MAP2 (1:1000) and GFAP (1:750) in normal antibody diluent overnight at 4°C. Next, cultures were washed three times with PBS-T and incubated with appropriate secondary antibodies (1:200 in normal antibody diluent) and DAPI (1:1000) for 30 min at room temperature. After incubation, cultures were washed twice with PBS-T and twice with PBS and imaged using a Keyence BZ-X-700 digital fluorescence microscope (Keyence, Itasca, IL). Images captured at 10× magnification were analyzed using Keyence BZ-X software to quantify the number of neurons, averaged across 25 fields/well, with 2–4 wells/treatment condition for each biological replicate.

For immunohistochemistry, brain tissues were fixed, paraffin-embedded, and sectioned at 5 µm thickness. Sections were incubated overnight at 55°C, followed by paraffin removal using Histoclear and rehydration with gradient ethanol. Next, sections were incubated with 3% hydrogen peroxide in methanol for 30 min to block endogenous peroxidase activity, followed by incubation with target retrieval solution (catalog #S169984-2, Thermo Fisher Scientific) for 30 min at 95°C for antigen retrieval. Sections were washed with PBS and blocked with 10% normal goat serum (catalog #5425, Cell Signaling Technology) for 60 min, followed by incubation with a primary rabbit antibody against p-eIF2α (1:1000) overnight at 4°C. Next, tyramide signal amplification (catalog #NEL700A001KT; Akoya Biosciences, Marlborough, MA) was performed using a FITC-conjugated goat anti-rabbit secondary antibody, per the manufacturer’s instructions. Next, sections were incubated with chicken MAP2 (1:200) and mouse GFAP (1:50) primary antibodies, followed by Cy3-conjugated goat-anti-chicken (1:200 dilution) and Cy5-conjugated goat-anti-mouse (1:200) secondary antibodies, respectively, before staining with DAPI (1:1000) to visualize nuclei.

### Measurement of interleukin 6

Astrocyte cultures plated in 200-µL medium in 24-well plates were treated with lipopolysaccharide (LPS from *E. coli.* O55:B5, catalog: L4524, Sigma), PA, or DMSO vehicle for 24 h. Supernatants were used to measure interleukin 6 (IL-6) with an enzyme-linked immunosorbent assay kit for mouse IL-6 (catalog #KMC0061; Thermo Fisher Scientific), according to the manufacturer’s protocol. Absorbance was quantified using a 96-well plate reader measuring at 450 nm.

### Quantitative real-time polymerase chain reaction

For quantitative real-time polymerase chain reaction (qPCR), after treatments, cultures and tissues were harvested in TRIzol reagent (catalog #15596018, Thermo Fisher Scientific), and RNA was extracted using the RNeasy lipid tissue mini kit (catalog #74804; Qiagen, Hildon, Germany), per manufacturers’ protocols. RNA quality was assessed using the A260/280 and A260/230 ratios using a nanodrop spectrophotometer (Thermo Fisher Scientific). The RT2 first strand cDNA synthesis kit (catalog #330404, Qiagen) with equal amounts of RNA was used to synthesize cDNA, which was diluted to 1:10. Gene expression levels were analyzed using TaqMan assays for *Hspa5* (BiP), *Ddit3* (Chop), *Ppp1r15a* (Gadd34), *Atf4* and *Eif2ak3* (PERK). *Gapdh* was used as the housekeeping control. The following TaqMan assays (catalog #4331182, Thermo Fisher Scientific) were used: *Gapdh* (Mm99999915_g1), *Hpsa5* (Mm00517691_m1), *Ddit3* (Mm01135937_g1), *Ppp1r15a* (Mm01205601_g1), *Atf4* (Mm00515325_g1), and *Eif2ak3* (Mm00438700_m1).

For TaqMan assays, cDNA was mixed with TaqMan reagent, TaqMan fast advanced master mix (catalog #4444557, Thermo Fisher Scientific), and water in 96-well qPCR plates and sealed with optical adhesive film. All samples were loaded in at least three technical replicates for qPCR analysis, with no-reserve transcription and no-template controls, and analyzed on a QuantStudio 3 device (Thermo Fisher Scientific) using the ÄÄCt method.

### Statistical analysis

For *in vivo* data, each mouse represented an independent biological replicate. For all cell culture experiments, enriched neuronal, neuroglial, and astrocyte cultures from each pregnant mouse constituted an independent biological replicate if the embryos/pups were dissected and pooled into a single culture before plating. Pups or embryos from the same dam but dissected separately were considered independent biological replicates. An untreated condition (UT) was also included within each biological replicate, and all results were normalized to the UT control. Unpaired Student’s *t* test, one-way analysis of variance (ANOVA) with repeated measures, or two-way ANOVA with Sidak’s multiple comparisons test were used as appropriate. Data were analyzed using GraphPad Prism statistical software (version 8.4.3) and presented as fold changes from UT ± standard error of the mean. All UT values were set to 1 or 100% and were not included in statistical analyses.

## Results

### PERK expression and activity in the brain are comparable between PERK-A/A and PERK-B/B mice

Accumulating evidence implicates altered PERK activity in neurodegenerative disorders^59,92–98^, suggesting PERK modulation as a potential therapeutic target^35,57–77,79–81,93^. In particular, the three exonic SNVs that are part of the PERK-B haplotype have been associated with increased risk for various disorders^85,99,100^, including neurodegenerative diseases such as progressive supranuclear palsy^52,53^, AD^51^, and HAND^53,84^. To investigate whether PERK-B modulated PERK expression levels or its activity in the CNS *in vivo*, we examined specific disease-relevant brain regions from PERK-A/A and the triple knock-in PERK-B/B mice^83^. Tissue homogenates from the cortex, hippocampus, striatum, and cerebellum of PERK-A/A and PERK-B/B mice were evaluated for PERK signaling by immunoblotting, revealing comparable levels of PERK expression in all brain regions between the PERK-A/A and PERK-B/B mice (Fig. 1A–E). Similarly, PERK activity, as determined with relative protein levels of phosphorylated eIF2α (p-eIF2α) and its downstream effector CHOP, were comparable in all regions between the PERK-A/A and PERK-B/B mice. Of note, the protein levels of BiP, were higher in the cortex and hippocampus of PERK-B/B mice than in those of PERK-A/A mice (Fig 1D). Immunohistochemical analysis of the cortex and hippocampus also indicated comparable p-eIF2α levels between PERK-A/A and PERK-B/B mice (Fig 1F-G).

**Figure 1:**
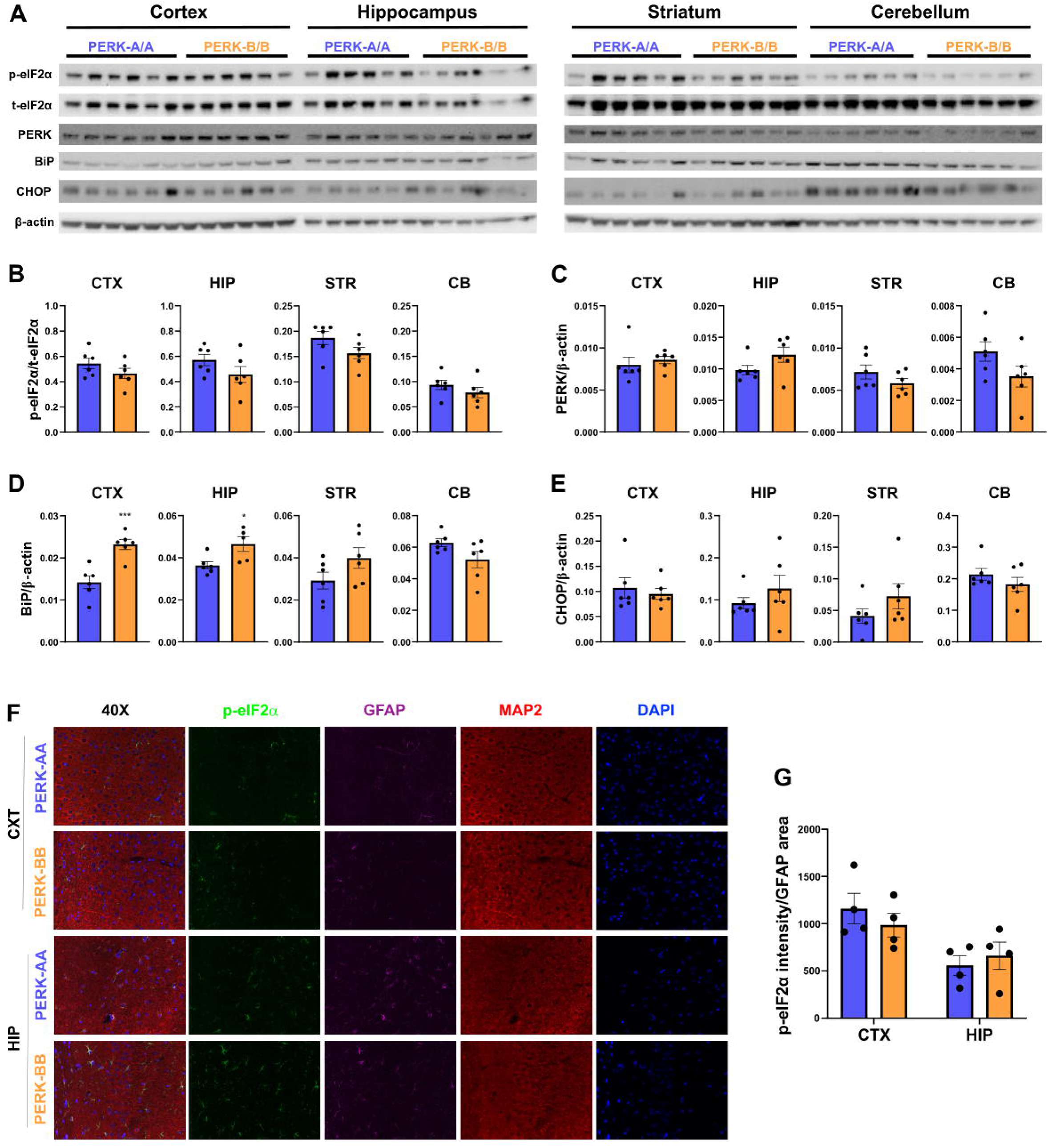
PERK expression and activity are comparable between PERK-A/A and PERK-B/B mice. **(A–E).** Tissue homogenates from cortex (CTX), hippocampus (HIP), striatum (STR), and cerebellum (CB) from PERK-A/A and PERK-B/B mice were prepared to determine the expression levels of p-eIF2α, t-eIF2α, PERK, BiP, CHOP, and β-actin with immunoblotting. Representative immunoblots (**A**) and quantification of p-eIF2α normalized to t-eIF2α, and PERK, BiP, and CHOP normalized to β-actin **(B–E). (F-G).** Representative 40× images (**F**) of hippocampal (HIP), and cortex (CTX) sections from PERK-A/A and PERK-B/B mice were stained for p-eIF2α, GFAP, MAP2, and DAPI using immunofluorescence and quantified for p-eIF2α intensity normalized to GFAP area **(F-G)**. Unpaired t-test; * p < 0.05, *** p < 0.001, N=5-6.

### PERK activation kinetics are different between PERK-A/A and PERK-B/B neuronal cultures

PERK activation and eIF2α phosphorylation are rapid, with most studies focusing on early time points ranging from 15 min to 2h^82,86^. However, some studies have reported that PERK signaling can be oscillatory in specific cell types^15,101^. Therefore, we compared PERK activation kinetics between PERK-A/A and PERK-B/B neurons. We monitored PERK activity over a 24-h period following treatment with 3 µM thapsigargin by measuring p-eIF2α levels. We observed distinct and slightly higher p-eIF2α levels in PERK-B/B neurons compared to PERK-A/A neurons (Fig 2A, 2B), suggesting altered PERK activation kinetics in PERK-B/B neurons. Notably, p-eIF2α levels did not significantly differ between the two groups after 2h of Tg treatment (Fig. 2C), which is the typical peak PERK activity in these cultures^94,102,103^. However, we detected significant differences in p-eIF2α levels at 6 h (Fig 2D), indicating altered PERK activation kinetics, with prolonged eIF2α phosphorylation in PERK-B/B cultures compared with PERK-A/A cultures.

**Figure 2:**
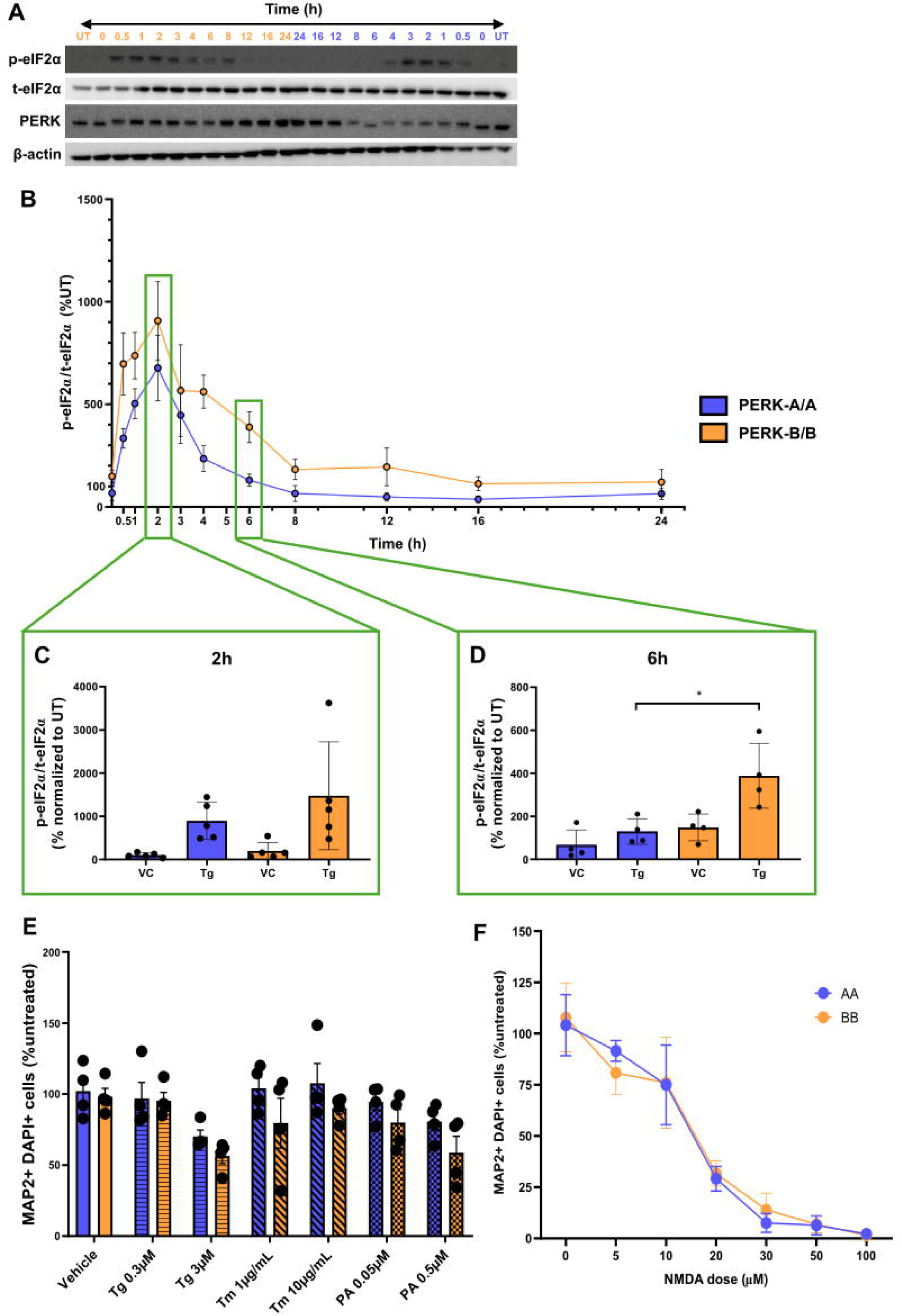
PERK-B/B neuronal cultures exhibit altered PERK activation kinetics compared to PERK-A/A cultures. **(A–D).** Cell lysates from PERK-A/A and PERK-B/B neuronal cultures treated with 3 µM thapsigargin for up to 24 h were analyzed with immunoblotting for p-eIF2α, t-eIF2α, PERK, and β-actin protein levels. Representative images **(A)** and quantifications of the p-eIF2α/t-eIF2α ratio **(B)** over the 24-h thapsigargin treatment time course, and quantification of the p-eIF2α/t-eIF2α ratio at 2 h **(C)** and 6 h **(D)** post thapsigargin treatment. n = 4–5. **(E-F)** DIV 18–21 neurons isolated from PERK-A/A or PERK-B/B mice were treated with indicated doses of thapsigargin (Tg), tunicamycin (Tm), or PERK-activator (PA) **(E)**, or NMDA **(F),** followed by immunofluorescence analysis to determine MAP2+/DAPI+ cells as a measure of surviving neurons, expressed as percentage change compared to untreated control. Two-way repeated-measures analysis of variance (ANOVA) with Sidak’s multiple comparisons comparing between genotypes for each treatment; n= 2-4; * p < 0.05.

We also determined whether PERK-B/B neurons exhibited alterations in other downstream signaling targets of PERK by measuring mRNA levels of *Atf4*, *Hspa5* (encoding BiP), *Ddit3* (encoding CHOP), and *Ppp1r15a* (encoding GADD34), in addition to *Eif2ak3*. The baseline expression levels did not significantly differ between PERK-A/A and PERK-B/B neurons for any of the interrogated targets (Supplemental Fig. 1A) at most time points following treatment with 3 µM thapsigargin (Supplemental Fig. 1B).

To investigate the functional implications of the observed changes in PERK signaling kinetics, we exposed primary neuronal cultures from PERK-A/A and PERK-B/B mice to low and high doses of various PERK inducers, including 0.3 and 3μM thapsigargin (Tg), which blocks ER calcium import, 1 and 10 μg/mL tunicamycin (Tm), which blocks N-linked glycosylation, and 0.05 and 0.5 μM of a synthetic PERK activator (PA) for 24 h. As shown in Fig. 2E, neuronal vulnerability to the ER stressors was comparable between PERK-A/A and PERK-B/B primary culture. We also used NMDA since excitotoxicity is implicated in the pathogenesis of neurodegeneration^104^. Similarly, treatment with increasing doses of NMDA (5–100 µM), which induces excitotoxicity^105^, did not lead to differences in neuronal viability between PERK-A/A and PERK-B/B neurons (Fig. 2F). These results suggested that, despite altered activation kinetics, PERK-B did not confer increased vulnerability to acute neuronal stressor exposure *in vitro*. This differed from our previous findings utilizing these mouse lines, where we found increased ER stress vulnerability in the liver and pancreas of PERK-B/B mice^83^.

### PERK-B/B-dependent differences PERK activation kinetics are lost in pure neuronal cultures

In the neuronal cultures above, AraC, a DNA damage inducer which is often used in primary neuron cell culture to remove astrocyte contamination, was not utilized to avoid off-target and neurotoxic effects^106^. However, prior research underscores the role of astrocytes in ER stress^38,107,108^, particularly the induction of inflammatory signaling cascades during PERK-mediated UPR^39^. Furthermore, immunohistochemical staining revealed that p-eIF2α staining in the brain was primarily localized to GFAP-positive cells (Fig. 1F-G), and we observed that PERK expression was significantly higher in astrocyte cultures as compared to neuron cultures (Supplementary Figure 3A-B). Therefore, we hypothesized that signaling changes seen in neuronal cultures may be driven by astrocytes, and the extent and effect of these changes is perhaps masked by in the mixed cultures. To confirm this, we repeated the assessment of PERK activation kinetics in neuronal cultures devoid of astrocytes using AraC treatment. Interestingly, we did not detect any significant differences in p-eIF2α levels over a 24-h period following treatment with 3 µM thapsigargin between PERK-A/A and B/B pure neuronal cultures (Fig. 3A–3D), suggesting that astrocytes present in neuronal cultures might account for the differences in activation kinetics between PERK-A/A and PERK-B/B cultures.

**Figure 3.**
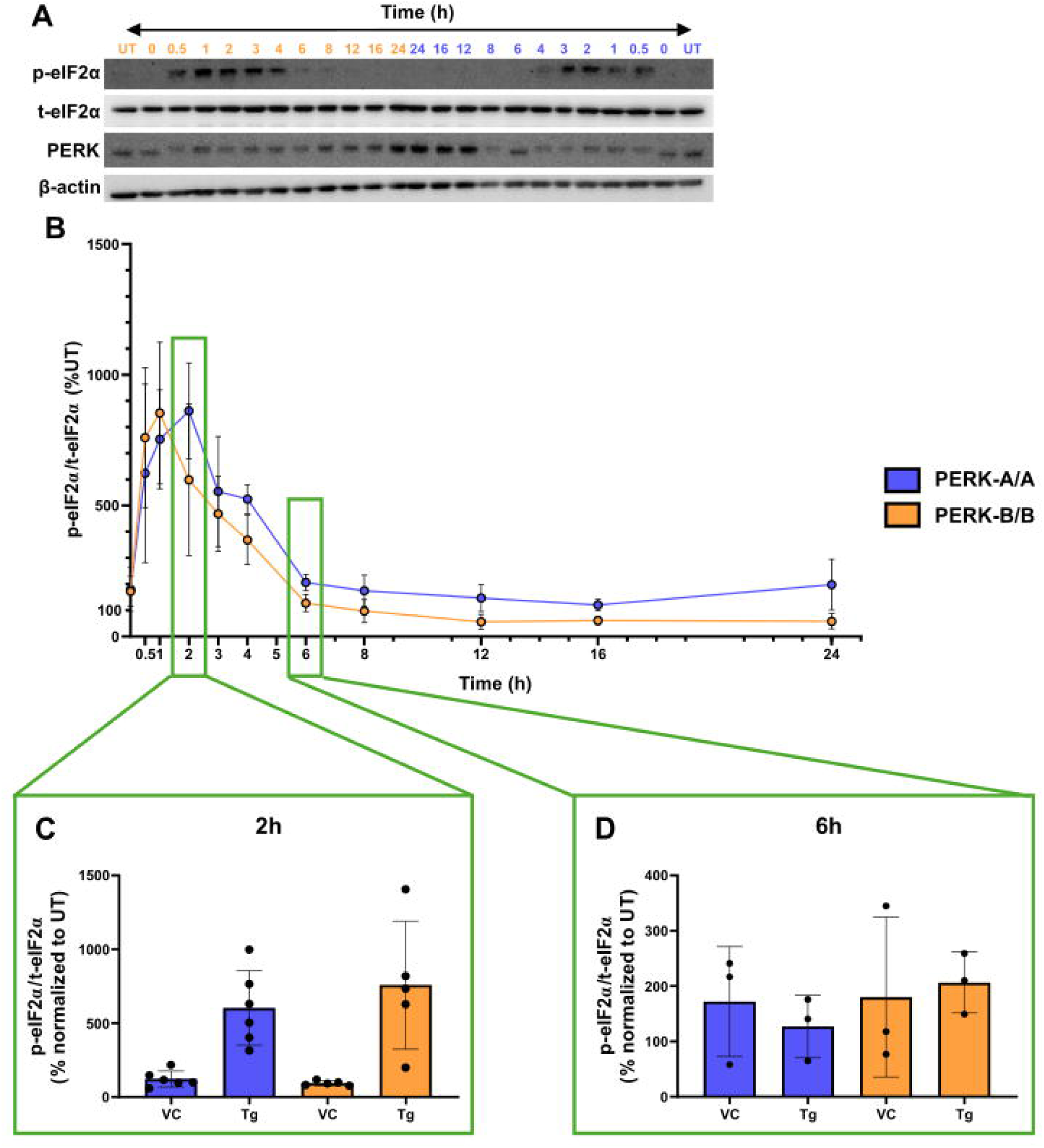
PERK activation kinetics are not different between PERK-A/A and PERK-B/B neuron-only cultures. **(A–D).** Cell lysates from PERK-A/A and PERK-B/B neuron-only cultures treated with 3 µM thapsigargin for up to 24 h were analyzed with immublotting for and p-eIF2α, t-eIF2α, PERK and β-actin protein levels. Representative blots **(A)** and quantifications **(B)** over the 24-h thapsigargin treatment time course, and quantification of the p-eIF2α/t-eIF2α ratio at 2 h **(C)** and 6 h **(D)** post thapsigargin treatment.. n = 3–6. Two-way repeated-measures ANOVA with Sidak’s multiple comparisons comparing between genotypes for each treatment.

### PERK activity is increased in PERK-B/B compared to PERK-A/A astrocytes cultures

Following discovery that acute PERK activity kinetics in neuronal cultures were lost in neuron-only cultures, discovery that PERK activation kinetics in neuronal cultures were lost in neuron only cultures, we compared primary astrocyte cultures from PERK-A/A and PERK-B/B mice. As shown in Fig 4A,B, PERK activity, assessed via p-eIF2α levels, was higher in PERK-B/B astrocytes than in PERK-A/A astrocytes following 2-h treatment with 0.3 µM thapsigargin. Interestingly, this difference was not observed when cultures were treated with 3 µM thapsigargin, suggesting that PERK-B/B might lower the activation threshold of PERK in primary astrocytes. Of note, in the same experimental paradigm, we did not observe any differences in PERK itself, nor downstream signals, between the PERK-A/A and PERK-B/B astrocytes at baseline (Supplemental Fig. 2A) or throughout the course of treatment with thapsigargin (Supplemental Fig. 2B), similar to neuronal cultures (Supplemental Figure 1).

**Figure 4:**
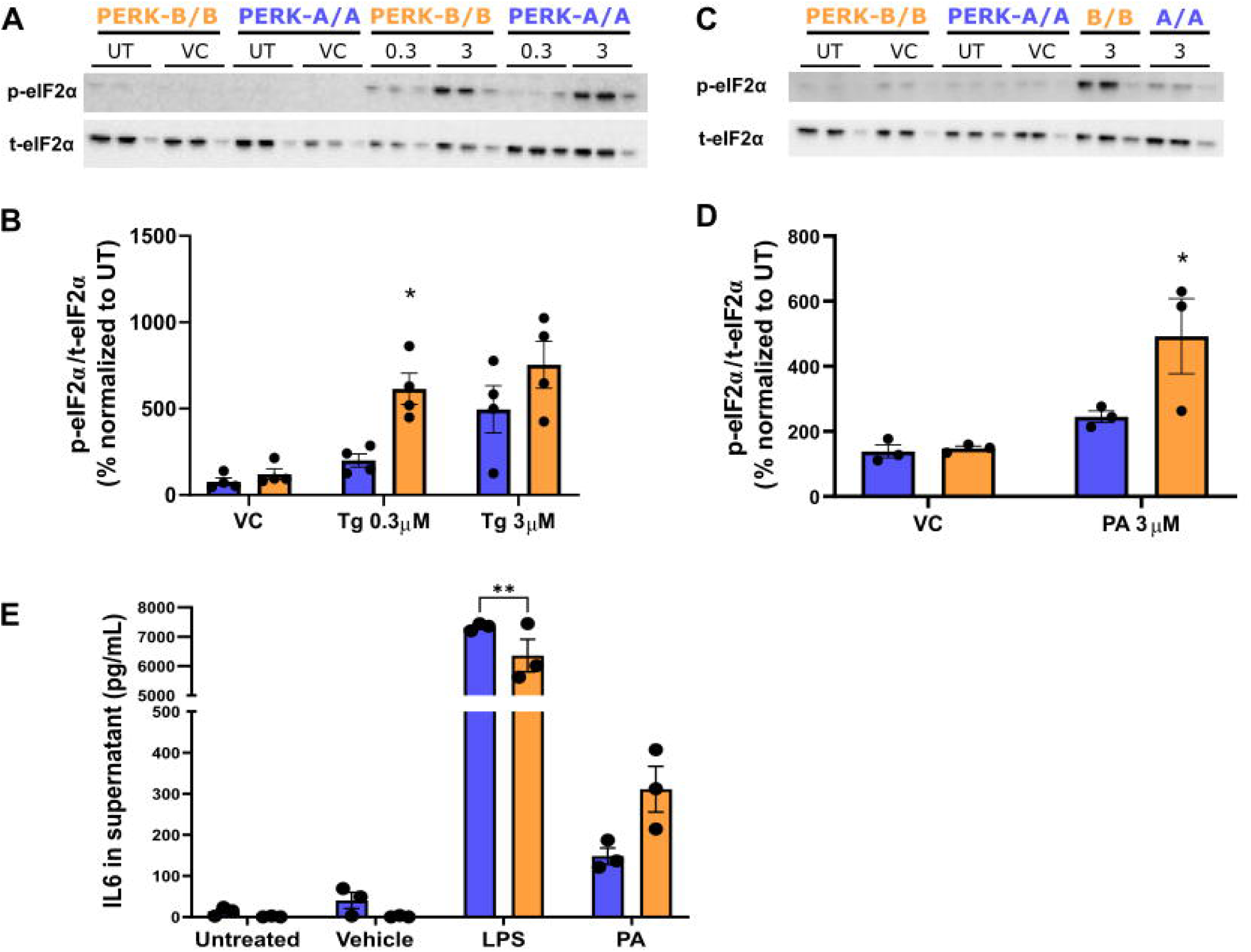
PERK activity is higher in PERK-B/B than PERK-A/A astrocyte cultures. **(A–D).** Cell lysates from PERK-A/A and PERK-B/B astrocyte cultures treated with thapsigargin (Tg) **(A-B)** or PERK activator (PA) **(C-D)** for 2h at the indicated doses were analyzed with immunoblotting for p-eIF2α and t-eIF2α protein levels. Representative blots **(A, C)** and quantification of the p-eIF2α/t-eIF2α ratio represented as percent change from untreated **(B, D).** n = 3–4. **(E).** Levels of IL-6, measured via enzyme-linked immunosorbent assay, in astrocyte supernatants 24 h after treatment with vehicle, PA, or LPS (positive control). Data are represented as raw values (pg/mL). Two-way repeated-measures ANOVA with Sidak’s multiple comparisons comparing between genotypes for each treatment, *p < 0.05; **p < 0.01

To further validate these findings, we utilized PA, which specifically targets and activates PERK, and observed that the PERK-B/B astrocytes exhibited higher PERK activity compared to PERK-A/A astrocytes following treatment with 3 µM PA for 2 h (Fig. 4C, D).

Astrocytes secrete IL-6 in response to ER stress, particularly downstream of PERK^40,41^. This secreted IL-6 is a mediator of neuroinflammation at the neuroimmune axis in many neurodegenerative disease models^40,109^. Moreover, the three exonic SNVs of the PERK-B haplotype have been linked to higher IL-6 levels in the cerebrospinal fluid of individuals with HIV^54^. Therefore, we measured IL-6 secretion following PA treatment for 24h in PERK-B/B and PERK-A/A astrocytes, and observed differences in IL-6 secretion (Fig. 4E). These results suggest that PERK-B/B alters PERK activity and ER stress response in astrocytes, potentially contributing to altered inflammatory responses.

### Acquired stress tolerance is impaired in PERK-B/B neuronal cultures

Exposure to low to mild levels of toxins or ER stressors can induce adaptive signaling that promotes survival and increases tolerance to a subsequent stressor^43,46–49^, whereas severe or prolonged ER stress can overwhelm these adaptive responses and promote cell death. This biphasic response is termed stress adaptation, hormesis, or preconditioning. A study in *C. elegans* emphasizes the central role of PERK in regulating ER stress-mediated hormesis^46^. To investigate whether the three SNVs knocked-in our PERK-B mice affect acquired stress tolerance in neurons, we utilized a stress tolerance paradigm, which included pretreating neuronal cultures with varying doses of PA (0.05 μM (low), 0.1 μM (medium) and 0.5 μM (high), which was then removed to allow recovery for 5 h, followed by a challenge with 10 μM NMDA treatment for 20 h (Fig. 5A). Following NMDA treatment, PERK-A/A neurons pretreated with low-dose PA exhibited increased survival compared to vehicle (DMSO) pretreatment (Fig 5B), indicating acquired stress tolerance to NMDA treatment, mediated by PA pretreatment. Of note, higher doses of PA or tunicamycin did not induce a protective effect. Importantly, PERK-B/B neurons did not exhibit acquired stress tolerance with any doses of PA. A similar genotype-dependent difference was observed following tunicamycin pretreatment, with PERK-A/A but not PERK-B/B cultures exhibiting acquired stress tolerance.

**Figure 5:**
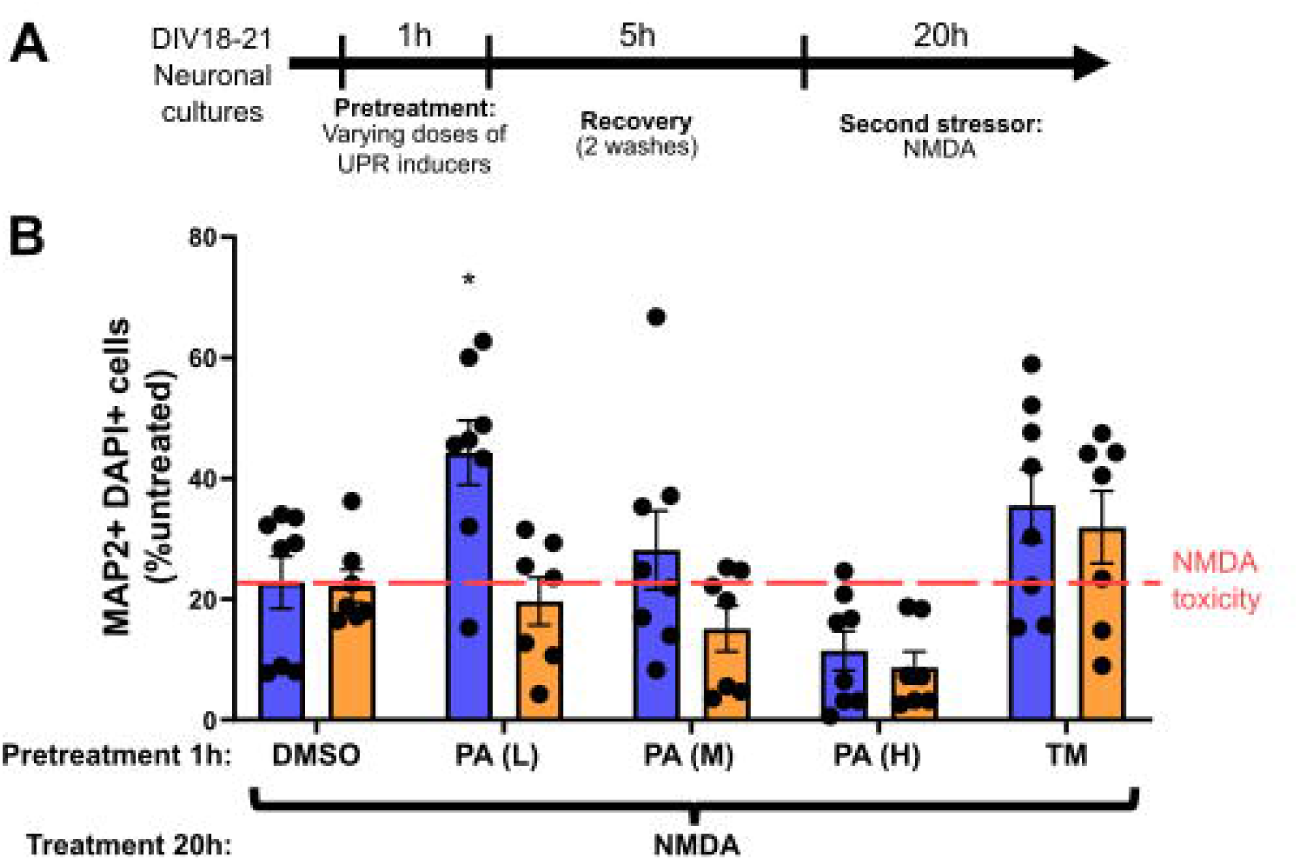
Acquired stress tolerance is impaired in PERK-B/B neuronal cultures. **(A–B).** Primary neuronal cultures obtained from PERK-A/A and PERK-B/B embryos were pretreated with either Tunicamycin (Tm) or a low (L) (0.05 μM), medium (M) (0.1 μM), or high (H) (0.5 μM) dose of PERK activator (PA) for 1h, followed by recovery for 5 h and subsequent treatment with 10 µM NMDA for 20 h **(A)**. Quantification of the number of MAP2+ DAPI+ cells across treatment groups expressed as percent change from untreated cultures **(B).** n = 7–8. Two-way repeated-measures ANOVA with Sidak’s multiple comparisons comparing between genotypes for each treatment; *p < 0.05.

### Impaired acquired stress tolerance in PERK-B/B neurons is mediated by astrocytes

To investigate whether astrocytes drove PERK-B-associated effects on acquired stress tolerance, we employed a neuron-astrocyte coculture model. We cultured pure neurons derived from PERK-A/A mice and co-cultured them with astrocytes obtained from PERK-A/A or Pz ERK-B/B mice (Fig. 6A). Using our stress adaptation paradigm, we observed a hormetic response in PERK-A/A neurons cocultured with PERK-A/A astrocytes, which exhibited improved survival in response to NMDA treatment following pretreatment with low-dose PA (Fig. 6B), mimicking our findings in neuronal cultures above (Figure 5). However, PERK-A/A neurons cocultured with PERK-B/B astrocytes did not exhibit a hormetic response (Fig. 6C), supporting an astrocyte-dependent impairment in acquired stress tolerance associated with PERK-B.

**Figure 6:**
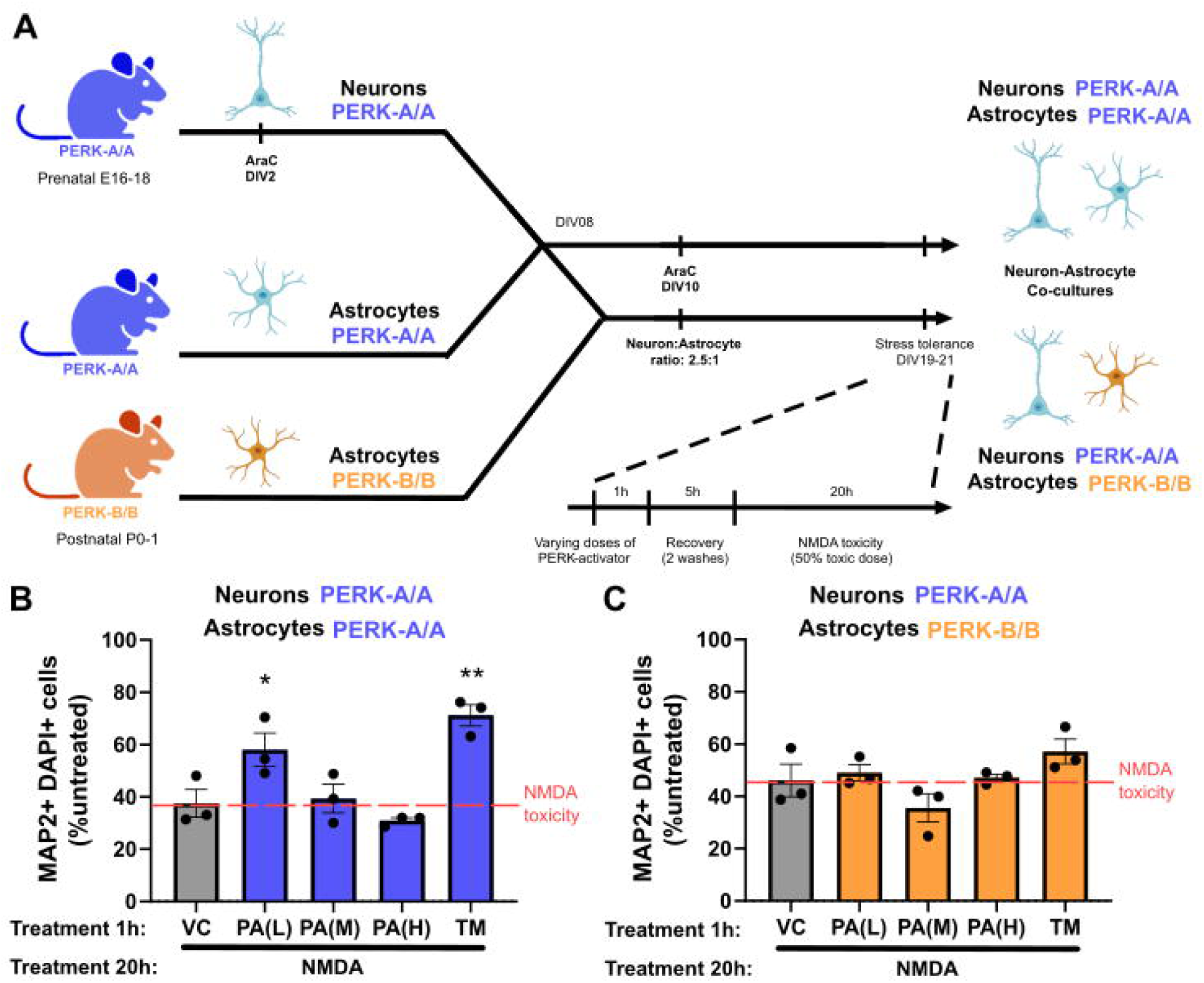
Impaired acquired stress tolerance in PERK-B/B neurons is mediated by astrocytes. **(A–C).** Astrocytes isolated from PERK-A/A or PERK-B/B mice were added to PERK-A/A neuronal cultures to generate astrocyte-neuron cocultures, which were pretreated with Tunicamycin (Tm) or varying doses (low (L) (0.05 μM), medium (M) (0.1 μM), or high (H) (0.5 μM)) of PERK activator (PA) for 1 h, followed by recovery for 5 h and subsequent treatment with 10 µM NMDA for 20 h **(A)**. Quantification of MAP2+DAPI+ cell counts normalized to untreated, for PERK-A/A neuron–PERK-A/A astrocyte cocultures **(B)** and PERK-A/A neuron–PERK-B/B astrocyte cocultures **(C)**. Data are presented as percent change from untreated. n = 3. One-way repeated-measures ANOVA with Dunnett’s multiple comparisons comparing each pretreatment to vechicle (DMSO); *p < 0.05, **p < 0.01.

## Discussion

We recently characterized the novel triple-knock-in PERK-B mice expressing the three exonic SNVs of the PERK B haplotype and demonstrated increased PERK activity associated with PERK-B *in vitro* and *in vivo* in the liver and pancreas of these animals^83^. In the present study we evaluated the effect of the exonic PERK-B SNVs in astrocytes and neurons and found that, while these SNVs caused minimal changes in most basal and acute PERK-associated stress markers, they did result in enhanced PERK activity in astrocytes, which appeared to contribute to prolonged PERK activity in astrocyte containing neuronal cultures, as well as impaired acquired stress tolerance in two-hit stress adaptation paradigms. The differences between PERK-A/A and PERK-B/B were evident in specific stress contexts and cell types, highlighting the subtle effects of the commonly occurring PERK-B haplotype and its potential contribution as a modifier of neurodegenerative disease risk in the presence of additional genetic or environmental factors.

Our findings contribute to accumulating evidence regarding the impact of PERK-B on PERK’s kinase activity, despite the conflicting data in previous *in vitro* studies^82,86,110^. PERK has been heavily implicated in neurodegeneration^55,56,59,92–98^ and is therefore considered a potential therapeutic target^35,57–77,79–81,93^. However, PERK signaling is far from linear; it exhibits oscillatory patterns within adaptable and redundant feedback networks^15,101^. Additionally, a plethora of pleiotropic downstream signaling pathways operate in concert based on the stress context, cell-specific regulation, and cellular and organismal homeostatic states. Our findings in this study provide additional context for the role of PERK-B and downstream signaling in stress response within specific CNS cell types.

First, we demonstrated that, at baseline, PERK kinase activity (determined by eIF2α phosphorylation) and PERK levels did not differ significantly between PERK-A/A and PERK-B/B mice in brain regions relevant to neurodegeneration, including the hippocampus, cortex, striatum, and cerebellum. Importantly, these observations were made in untreated mice, reflecting baseline PERK activity, and do not preclude potential differences in PERK activity and downstream signaling between PERK-A/A and PERK-B/B mice in response to ER stress *in vivo*. Conversely, BiP levels in the hippocampus and cortex were higher in PERK-B/B mice than in PERK-A/A mice, suggesting a compensatory mechanism that modulates PERK and normalizes its activity. Given that both the hippocampus and cortex exhibit significant neural activity and are likely to have higher cellular and local protein translation, ER stress might already be present at baseline in these regions compared to other brain regions. Furthermore, specific cell populations in these regions may be more responsive to subtle changes in PERK activity and ER stress. In fact, we have previously reported increased BiP levels in astrocytes and neurons, but not in macrophages/microglia, in postmortem cortical tissue from individuals with HAND^111^, underscoring BiP’s role as a marker of ER stress in neurodegeneration and emphasizing its cell-specific expression.

The immunohistochemical analysis confirmed our immunoblot results, showing that the baseline PERK activity in the hippocampus and cortex was comparable between PERK-A/A and PERK-B/B mice. Notably, by immunohistochemical staining, p-eIF2α expression was more pronounced in GFAP-positive astrocytes than in neurons. GFAP positivity supports astrocyte activation, highlighting the role of astrocytes in PERK-mediated UPR in the CNS and, again, the presence of ER stress at baseline in these regions.

In *in vitro* studies assessing the impact of PERK-B SNVs and the ER stress response on CNS-relevant cell populations, including neurons and astrocytes, we observed differences in PERK activation kinetics between PERK-A/A and PERK-B/B neurons. In both lines, peak kinase activity occurred within 2–3 h. While PERK activity began to normalize in PERK-A/A neurons thereafter, it remained active for another 4–6 h in PERK-B/B neurons, implicating that PERK-B might be associated with differential regulatory pathways, particularly those involving negative feedback mechanisms that govern PERK deactivation. PERK-B SNVs may also impact the interaction of PERK with PERK-specific phosphatases, the identities of which remain elusive. Furthermore, PERK-B SNVs may affect PERK’s binding affinity to BiP, as BiP upregulation during ER stress titrates unfolded proteins off PERK, leading to its deactivation.

While these results implicated differences between PERK-A/A and PERK-B/B neurons, the neuronal cultures used in these studies contain up to 10% astrocytes. Strikingly, the observed difference in PERK activation kinetics was not observed in pure neurons, underscoring the potential cell-type-specific effects of PERK-B. Notably, we observed higher PERK activity in PERK-B/B astrocytes treated with low doses of thapsigargin or PA. Astrocytes and neurons exhibit distinct responses to stress. Astrocytes are strongly activated under metabolic stress, inducing autophagy, whereas autophagy induction is relatively muted in neurons^112^. Paradoxically, neurons are more sensitive to inhibition of the ubiquitin-proteasome system and proteotoxicity^113^. Astrocytes play dual roles in response to oxidative stress: they provide neuroprotection against mild oxidative stress but can exacerbate oxidative stress under more severe stress, such as that encountered in pathologic conditions. In contrast, neurons exhibit lower reactivity but are more vulnerable to reactive oxygen species than astrocytes^114^. A study demonstrated that neurons were considerably more tolerant than astrocytes to perturbations in PERK signaling downstream of ER stress, due to redundant signaling pathways that ensure proteostatic resilience^115^. Interestingly, astrocytes uniquely express an additional ER stress sensor, old astrocyte specifically induced substance (OASIS) ^116,117^, a factor not expressed in neurons and implicated in the activation of inflammatory pathways^38^. Moreover, astrocytes contribute to cell-nonautonomous ER stress by transmitting ER stress signals via secreted soluble factors. These signals may lead to the secretion of IL-6 and IL-1β from microglia *in vitro*^42^ and precipitate neurodegeneration *in vivo*^6^. Additionally, under ER stress, astrocytes can secrete IL-6 in a PERK-dependent manner^41^. Our observation that stressed PERK-B/B astrocytes also exhibited increased IL-6 secretion compared to PERK-A/A astrocytes provides new insights into the role of PERK in inflammation and highlights the complex, cell-type- and stress-context-specific regulation of proteostasis and subsequent cellular pathways, including inflammation.

Previous studies have shown that PERK mediates ER stress-induced IL-6 secretion, potentially through the JAK1/STAT3 and the NFKB signaling cascades^39,41,118,119^. Importantly, we previously reported that PERK-B SNVs were associated with elevated IL-6 levels in the cerebrospinal fluid of individuals with HAND^54^. The implications of PERK-B-mediated changes in ER stress-induced inflammation remain unclear; however, we previously demonstrated higher PERK activity in mouse monocyte-derived macrophages isolated from PERK-B/B mice compared with those isolated from PERK-A/A mice^83^. Collectively, these findings suggest that the effects of PERK-B extend to astrocyte and myeloid cell functions, contributing to pathology in neurodegenerative diseases with an inflammatory component.

Surprisingly, neurotoxicity induced by ER stressors or NMDA was not different between PERK-A/A and PERK-B/B neurons, highlighting the nuanced effects of PERK-B that may not lead to overt toxicity or pathology in non-chronic models, or in the absence of additional genetic or environmental risk factors. Indeed, PERK-B was reportedly associated with increased risk of AD in a Han Chinese cohort of individuals harboring *APOE* ε*4*, the strongest genetic risk factor for AD^51^. Moreover, the majority of studies implicating PERK-B as a disease modifier were conducted in Asian cohorts^51,85,99,120^. Similarly, we previously reported the association of PERK-B SNVs with higher risk for neurocognitive impairment and depression in people with HIV who were on antiretroviral treatment^54,84^; HIV and antiretroviral drugs can initiate or precipitate ER stress^88,111,121,122^. The effect size of common genetic variants may be altered through specific gene–gene and gene–environment interactions^123^; therefore, these studies suggest that additional factors synergize with PERK-B to influence disease risk.

In addition to its role in survival under ER stress, we also investigated PERK’s impact on acquired stress tolerance. Hormesis represents a non-linear dose-response relationship with toxins or stressors, characterized by low-dose stimulation and high-dose inhibition. Low/mild doses of ER stress lead to adaptive changes, including expansion of the ER membrane, induction of BiP, and activation of prosurvival pathways that persist, resulting in resilience and enabling the resolution of subsequent stressors without commitment to apoptosis^43,47–50,124–127^. Hormesis is an evolutionary adaptation used by all life forms to cope with changing environmental conditions, especially the presence of toxins or stressors. Previous studies have implicated ER stress, specifically PERK, in the regulation of ER-stress-related hormetic effects^43,46^. In the present study, we utilized neuronal and astrocyte-neuron cocultures to demonstrate that PERK variant type influenced PERK regulation of ER stress-mediated hormetic effects. We used a ‘two-hit’ paradigm, pretreating cells with ER stressors, followed by a second challenge. Previous studies employing comparable models have assessed stress resilience *in vitro*^43,125–127^ and *in vivo*^44,45^ by exposing the experimental model to a second challenge specific to their paradigm, such as glutamate for hippocampal neurons and lipopolysaccharide for microglia and astrocytes. In the present study, we pretreated cultures with a PERK-specific activator, followed by exposure to an excitotoxic dose of NMDA, mimicking conditions akin to neurodegeneration^104^. Importantly, our findings suggest that PERK-B is associated with impaired acquired stress tolerance and that this effect is primarily driven by astrocytes. Other studies examining cell-type-specific effects of proteostatic and oxidative stress-mediated hormesis showed that astrocytes, unlike neurons, exhibited a robust hormetic effect^125–127^. Also utilizing a two-hit paradigm, these studies demonstrated that activated astrocytes, when cocultured with neurons, developed resilience and provided protection against additional stressors. Although the mechanisms underlying the loss of stress tolerance with PERK-B remain unclear, it is possible that altered stress response with PERK-B may contribute to these effects. The altered signaling kinetics observed with PERK-B during the acute stress-tolerance paradigm, suggest that differences in stress-response resolution may contribute to the impaired acquired stress. Understanding whether the diminished stress tolerance observed with PERK-B implies loss of function or dampening of prosurvival pathways, gain of toxic function or activated proapoptotic cascades, or a combination of these factors in distinct phases of stress response requires further investigation. These effects might also be attributed to PERK’s role in endogenous antioxidant response, which warrants further investigation^47^.

Hormesis typically follows a bell-shaped curve^47,48^, where low/mild doses of ER stressors yield beneficial effects within a hormetic window. At the other extreme, higher doses of ER stressors exacerbate cell survival under additional stressors, leading to reduced resilience. We suggest that this window might shift in PERK-B-expressing cells such as astrocytes, given that its effects are mainly observed in mild stress conditions. Elucidating the definition and determinants of the hormetic window *in vivo* is crucial for the development of adjunctive therapies for individuals harboring PERK-B. This window may be influenced by other genetic and environmental factors, including age, ancestry, sex, and the presence of other genetic disease risk modifiers, such as *APOE* ε*4* in neurodegeneration. Understanding these nuances is essential for patient-tailored therapeutic strategies.

In conclusion, we investigated the effects of PERK-B SNVs on stress response and stress tolerance in CNS-relevant cell populations derived from PERK-A/A or PERK-B/B mice. While PERK-B/B mice did not exhibit significantly altered PERK levels or kinase activity *in vivo*, it was associated with elevated BiP levels in the hippocampus and cortex of PERK-B/B mice. *In vitro,* PERK-B/B neurons exhibited subtle alterations in activation kinetics in the form of prolonged activation compared to PERK-A/A. Surprisingly, both PERK-A/A and PERK-B/B neurons exhibited comparable neurotoxicity with various ER stress inducers, suggesting that the subtle effects of PERK-B might be unmasked only in the presence of additional factors. Notably, stress tolerance was lower in PERK/B/B neurons and neuron-astrocyte cocultures *in vitro*, an effect driven by astrocytes, indicating that PERK-B/B cells were less resilient than PERK-A and exhibited altered hormetic response. These findings shed light on the nuanced impact of PERK-B in specific stress contexts and cell types, potentially enhancing the genetic risk for neurodegeneration in conjunction with other factors. Further research is necessary to fully comprehend the underlying molecular mechanisms and to determine if PERK-B acts as a gain of toxic function or loss of physiologic function.

