## Supplemental Figures for "Genetic variation in *Eif2ak3* alters astrocyte-dependent stress tolerance in neurons"

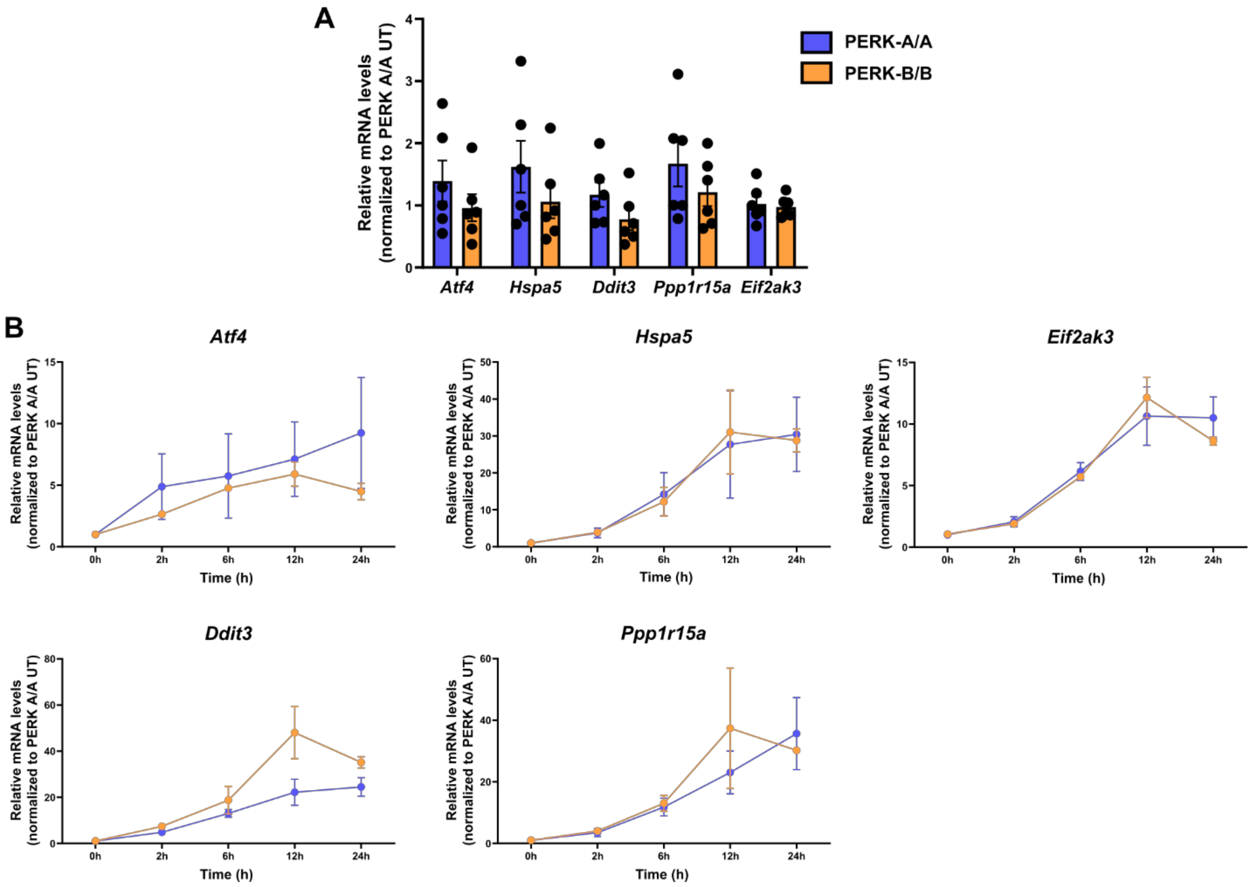


**Supplemental Figure 1: Downstream PERK signaling is comparable between PERK-A/A and PERK-B/B neuronal cultures.** cDNA was synthesized from RNA collected from PERK-A/A and PERK-B/B neuronal cultures treated with 3 μM thapsigargin for 0-24h. mRNA expression for the genes coding for ATF4 (*Atf4)*, Bip (*Hspa5*), CHOP (*Ddit3*), GADD34 (*Ppp1r15a*), PERK (*Eif2ak3*) and GAPDH (*Gapdh*) was assessed using qPCR. Relative gene expression of *Atf4*, *Hspa5*, *Ddit3*, *Ppp1r15a* and *Eif2ak3*, normalized to *Gapdh* at baseline **(A)** and at all post-treatment time points **(B)**. All data represented as a percentage of untreated. Two-way ANOVA with Sidak's multiple comparisons with repeated measures, n=2-6.

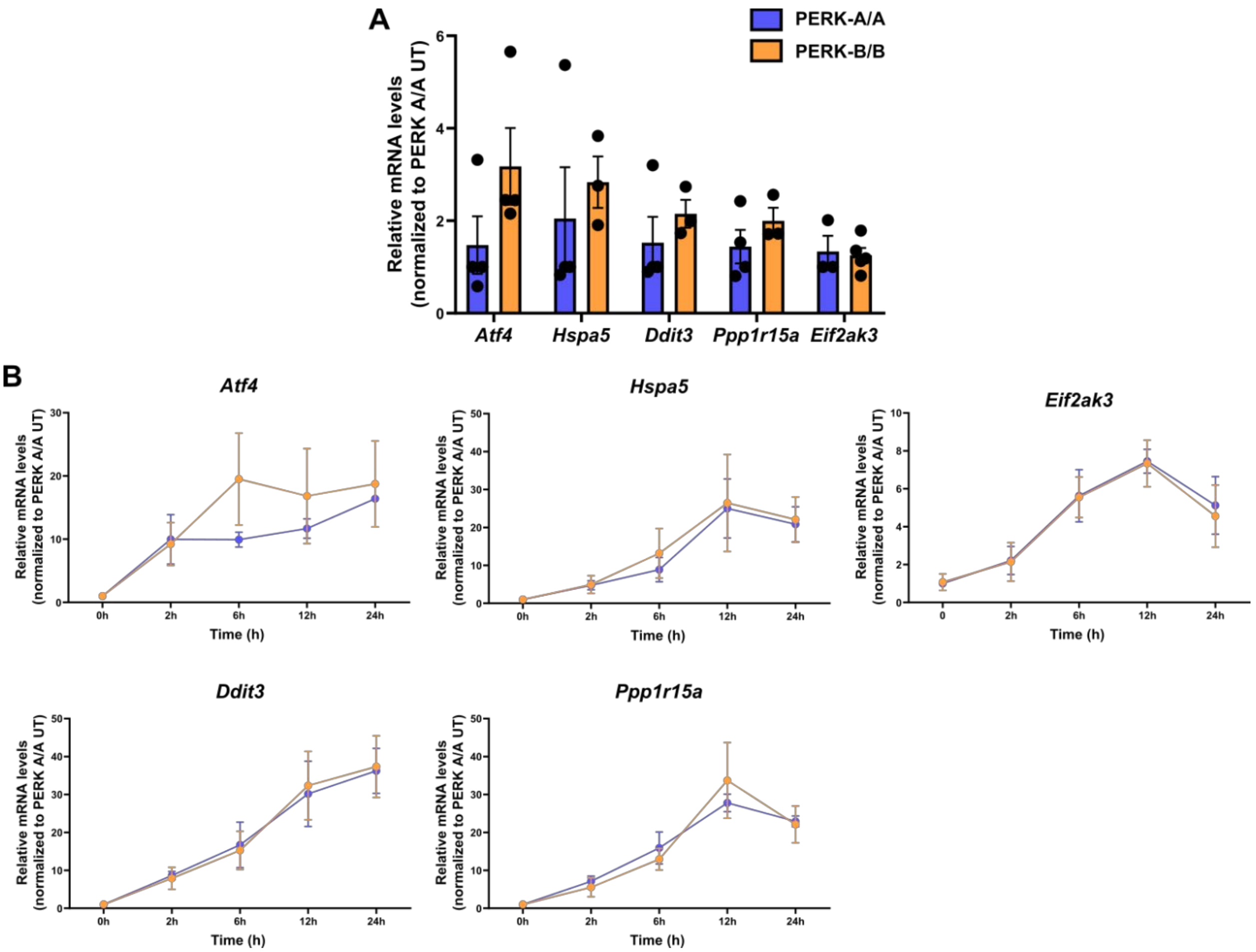


**Supplemental Figure 2: Downstream PERK signaling is comparable between PERK-A/A and PERK-B/B astrocytes.** cDNA was synthesized from RNA collected from PERK-A/A and PERK-B/B astrocytes treated with either 3 µM thapsigargin for 0-24h. mRNA expression for the genes coding for ATF4 (*Atf4)*, Bip (*Hspa5*), CHOP (*Ddit3*), GADD34 (*Ppp1r15a*), PERK (*Eif2ak3*) and GAPDH (*Gapdh*) was assessed using qPCR. Relative gene expression of *Atf4*, *Hspa5*, *Ddit3*, *Ppp1r15a* and *Eif2ak3*, normalized to *Gapdh* at baseline **(A)** and at all post-treatment time points **(B)**. All data represented as a percentage of untreated. Two-way ANOVA with Sidak's multiple comparisons with repeated measures, n=3-5.

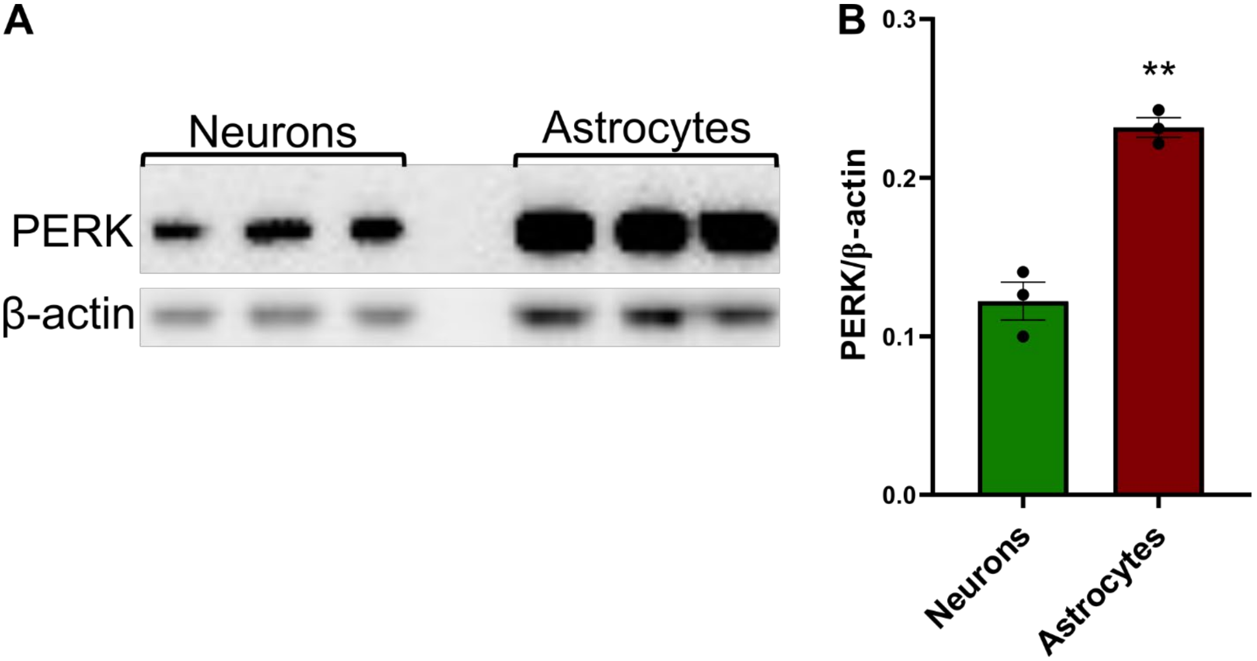


**Supplemental Figure 3: PERK protein levels are higher in astrocyte cultures compared to neuronal cultures.** Cell lysates from PERK-A/A neuronal and astrocyte cultures were analyzed with immunoblotting for PERK and β-actin protein levels. Representative blots **(A)** and quantifications of PERK protein levels normalized to β-actin **(B)**. n = 3. Unpaired t-test; ** $p < 0.01$.
